# A LAMP at the end of the tunnel: a rapid, field deployable assay for the kauri dieback pathogen, *Phytophthora agathidicida*

**DOI:** 10.1101/793844

**Authors:** Richard C. Winkworth, Briana C.W. Nelson, Stanley E. Bellgard, Chantal M. Probst, Patricia A. McLenachan, Peter J. Lockhart

## Abstract

The collar rot causing oomycete, *Phytophthora agathidicida*, threatens the long-term survival of the iconic New Zealand kauri. Currently, testing for this pathogen involves an extended soil bioassay that takes 14-20 days and requires specialised staff, consumables, and infrastructure. Here we describe a loop-mediated isothermal amplification (LAMP) assay for the detection of *P. agathidicida* that targets a portion of the mitochondrial apocytochrome b coding sequence. This assay has high specificity and sensitivity; it did not cross react with a range of other *Phytophthora* isolates and detected as little as 1 fg of total *P. agathidicida* DNA or 116 copies of the target locus. Assay performance was further investigated by testing plant tissue baits from flooded soil samples using both the extended bioassay and LAMP testing of DNA extracted from baits. In these comparisons, *P. agathidicida* was detected more frequently using the LAMP assay. In addition to greater sensitivity, by removing the need for culturing, the hybrid baiting plus LAMP approach is more cost effective than the bioassay and, importantly, does not require a centralised laboratory facility with specialised staff, consumables, and equipment. Such testing will allow us to address outstanding questions about *P. agathidicida*. For example, the hybrid approach could enable monitoring of the pathogen beyond areas with visible disease symptoms, allow direct evaluation of rates and patterns of spread, and allow the effectiveness of disease control to be evaluated. The hybrid assay also has the potential to empower local communities. These communities could use this diagnostic tool to evaluate the pathogen status of local kauri stands, providing information around which to base their management and allowing informed engagement with wider initiatives.

## Introduction

The long-term survival of kauri, *Agathis australis* (D.Don) Loudon (Araucariaceae), is threatened by the oomycete *Phytophthora agathidicida* B.S. Weir, Beever, Pennycook & Bellgard (Peronosporaceae) [1]. This soil-borne pathogen causes a collar rot that results in yellowing of the foliage, bleeding cankers on the lower trunk, thinning of the canopy and eventually in tree death. Initially reported from Great Barrier Island in the early 1970’s [2], since the late 1990’s the disease has spread rapidly across northern New Zealand [3].

Broad phylogenies for the Peronosporaceae (e.g., 4-7) have identified strongly supported sub-clades. For example, Bourret et al. [7] described total of 16 sub-clades, the majority of which contain at least one *Phytophthora* species. The phylogenetic analysis of Weir et al. [8] placed *P. agathidicida*, along with three other formally recognized species, within Clade 5. Although *P. agathidicida* is currently the only clade 5 species reported from New Zealand, it is not the only *Phytophthora* species associated with kauri forests. A further five *Phytophthora* species have been reported from kauri forest soils. Specifically, *P. chlamydospora* Brasier and Hansen (Clade 6), *P. cinnamomi* Rands (Clade 7), *P. cryptogea* Pethybr. & Laff. (Clade 8), *P. kernoviae* Brasier (Clade 10), and *P. nicotianae* Breda de Haan (Clade 1) [1, 3]. Other oomycetes, including several *Pythium* (Pythiaceae) species, have also been reported from kauri forest soils [3].

Methods for the isolation and identification of *Phytophthora* typically involve recovery from infected host root material [9, 10] or from soil samples using plant tissue fragments (e.g., detached leaves) as bait for zoospores [11, 12]. Currently, the preferred method of testing for *P. agathidicida* is an extended soil bioassay [13]. Briefly, soil samples are first air dried (2-8 days), then moist incubated (4 days), and finally flooded with water and baited (2 days). At this point the baits are surface sterilised, plated onto *Phytophthora*-selective media, and incubated at 18°C (6 days). *Phytophthora agathidicida* is identified from the resulting cultures using morphology or molecular diagnostics both of which require trained laboratory staff. The former is typically based upon the size and reflection of the antheridium [14] while an RT-PCR assay targeting the internal transcribed spacer (ITS) regions of the ribosomal DNA [15] is routinely used for identification of cultured *P. agathidicida* isolates.

Increasing the immediacy of results from diagnostic testing can substantially improve outcomes in terms of disease control. Traditionally, shortening the time needed to diagnose a disease has involved the use of new protocols or tools in the laboratory. For example, replacing culture-based methods with molecular diagnostics. However, the use of approaches that can be conducted onsite, thereby reducing reliance on centralised laboratory facilities is becoming an increasingly important means of enhancing the immediacy of results [16, 17]. The emergence of new amplification technologies is enabling the development of rapid, field-deployable approaches to genetic diagnostics [18]. In particular, loop-mediated isothermal amplification (LAMP), an approach that combines rapid amplification with high specificity and sensitivity, is rapidly becoming an important diagnostic tool, especially for point of care applications. There are now numerous examples of LAMP tests from fields including human medicine [e.g., 19, 20], agriculture [e.g., 21, 22], and animal health [e.g., 23, 24]. LAMP has several features that make it particularly well suited to non-laboratory applications. First, the DNA polymerases used to catalyse the LAMP reaction have strand-displacement activity. Therefore, unlike PCR-based diagnostics, LAMP assays may be carried out using relatively simple equipment [25]. Second, these polymerases are less sensitive to inhibitors than those used in PCR reactions. This allows simpler methods for DNA isolation to be used [26]. Finally, turbidity or colorimetry can be used for end-point detection of LAMP products further reducing reliance on sophisticated equipment [26].

The cost of the extended soil bioassay together with the requirement for specialised staff and infrastructure means that, typically, this assay is conducted only after the appearance of physical disease symptoms. Such restricted testing severely limits our basic understanding of *P. agathidicida* and our ability to make informed management decisions at local, regional and national scales. For example, we are yet to characterise the distribution of the pathogen, how fast it is spreading, or the efficacy of interventions (e.g., track closures) aimed at disease control. Addressing these knowledge gaps requires ongoing, active monitoring of both diseased and healthy sites across the distribution of kauri. This cannot be achieved using the existing test. Instead, a reliable and rapid assay for *P. agathidicida* that is both cost effective and robust enough to be deployed outside of a laboratory is needed. Beyond increasing existing capacity, such testing has the potential to enable individual landowners and community groups to evaluate pathogen status in their area and thereby engage in an informed way with regional and national initiatives.

LAMP assays have already been reported for the detection of several *Phytophthora* species. These include tests for *P. capsici* Leonian [27], *P. cinnamomi* [28], *P. infestans* (Mont.) de Bary [29, 30], *P. melonis* Katsura [31], *P. nicotinae* [32], *P. ramorum* Werres, De Cock & Man in ‘t Veld [22] and *P. sojae* Kaufm. & Gerd. [33]. The genetic targets of these tests include the Ras-related protein (*ypt1*) gene [i.e., 30-33] and the ITS regions [i.e., 22, 27]. As might be expected, LAMP tests for *Phytophthora* species differ in terms of both their absolute detection limits and performance relative to PCR-based assays. For example, Hansen et al. [29] and Khan et al. [30] reported *P. infestans* LAMP tests with detection limits of between 128 fg and 200 pg. The most sensitive of these, that of Khan et al. [30], was ten times more sensitive than a test based on nested PCR and at least 100 times more sensitive than either RT-PCR or conventional PCR tests for the corresponding locus.

Here we describe a hybrid bioassay for the detection of *P. agathidicida*. This combines conventional soil baiting with a highly specific and sensitive LAMP assay to directly test the plant bait tissues for the presence of the pathogen. By reducing assay cost, the time needed for pathogen detection, and reliance on centralised laboratories this approach has the potential both to overcome key limitations of the currently used extended soil bioassay and provide data that will inform our basic understanding of the disease and its management.

## Materials and Methods

### Target region identification

To identify potential targets for genetic testing, publically available *Phytophthora* and *Pythium* mitochondrial genome sequences were obtained from the NCBI RefSeq database (https://www.ncbi.nlm.nih.gov). These were combined with mitochondrial genome sequences from *Phytophthora* and related taxa (e.g., *Pythium, Plasmopara*) assembled at Massey University (Winkworth et al., unpubl.). The combined collection comprised mitochondrial genomes from 25 species representing 12 of the 16 clades of *Phytophthora* and downy mildews reported by Bourret et al. [7]. This sample included all five *Phytophthora* species reported from kauri forests [1, 3], all currently recognized representatives of *Phytophthora* clade 5 [8], and two accessions of *P. agathidicida* [8]. A list of species and accession numbers for publicly available sequence data are provided in S1 Table.

Comparisons of whole mitochondrial genome sequences for recognized *Phytophthora* clade 5 species have shown these genomes to be co-linear (Winkworth et al., in prep). To identify potential targets for LAMP assays, mitochondrial genome sequences from five *Phytophthora* clade 5 species were aligned using the MUSCLE [34] alignment tool as implemented in Geneious V9.1.2 [35]. To evaluate the utility of the identified loci, 0.5-1 kb sections of DNA sequence containing these regions were extracted from all 25 mitochondrial genomes and multiple sequence alignments for each constructed as before.

### LAMP primer design

LAMP primer sets were generated for potential target regions using PrimerExplorer v5 (https://primerexplorer.jp/e/). For each region, an initial search for regular primers used the *P. agathidicida* sequence along with default parameter values; the GC content threshold was progressively lowered in subsequent searches until at least one regular primer set was recovered. Candidate primer sets were then compared to multiple sequence alignments for the corresponding target locus; primer sets where annealing sites did not distinguish *P. agathidicida* were not considered further. For the remaining primer sets, a search was then made for loop primers using PrimerExplorer v5 and the same GC content threshold as generated the regular primer set.

Finally, each primer set was queried against the NCBI nucleotide database (https://www.ncbi.nlm.nih.gov) using BLAST [36] to investigate non-specific annealing.

### LAMP assay optimisation

For reaction optimisation, all LAMP assays were conducted in 25 μL volumes consisting of 15 μL Optigene Isothermal Master Mix (OptiGene Ltd., Horsham, West Sussex, England) plus primer cocktail, extracted DNA and milliQ water (H_2_O). Volumes of the three latter components were varied depending on the reaction conditions and template concentration. Reaction sets typically included both positive (i.e., 2 ng total DNA from cultured *P. agathidicida* isolates NZFS 3128 or ICMP 18244; see S1 Table for accession details) and negative (i.e., no DNA) controls. LAMP assays were performed using a BioRanger LAMP device (Diagenetix, Inc., Honolulu, HI).

Initial optimisation of the LAMP assay evaluated three parameters. First, we investigated the impact of varying the ratio of F3/B3 to FIP/BIP primer pairs. Ratios of F3/B3 to FIP/BIP primers of 1:3, 1:4, 1:6, and 1:8 were trialled; in each case the final concentration of the F3 and B3 primers was maintained at 0.2 μM with the concentrations of FIP and BIP primers being 0.6 μM, 0.8 μM, 1.2 μM, and 1.6 μM, respectively. Second, we examined the effect of amplification temperature. LAMP assays were performed at amplification temperatures of 60°C, 63°C, and 65°C. In all cases, DNA amplification was followed by enzyme denaturation at 80°C for 5 min. Finally, the amplification time was varied from 30-90 minutes.

### LAMP assay specificity

The specificity of the *P. agathidicida* LAMP assay was evaluated using optimised reaction mixes and conditions. These tests were conducted using 2 pg total DNA from six *P. agathidicida* isolates as well as from isolates of 11 other *Phytophthora* species representing nine of the 16 clades reported by Bourret *et al*. [7]. All *Phytophthora* species reported from kauri forests and all currently recognized representatives of *Phytophthora* clade 5 were included in the test set (S1 Table). Individual reaction sets also included both positive (i.e., 2 ng total DNA from cultured *P. agathidicida* isolates NZFS 3128 or ICMP 18244) and negative (i.e., no DNA) controls.

### LAMP assay sensitivity

We first evaluated LAMP assay sensitivity in terms of total *P. agathidicida* DNA. For these tests, a ten-fold dilution series with between 1 ng and 10 ag of total DNA from cultured *P. agathidicida* isolate ICMP 18244 together with optimised reaction mixes and conditions were used. Testing of the dilution series was conducted both with and without the addition of DNA from a plant tissue commonly used when baiting for *P. agathidicida*; specifically, 2 ng total *Cedrus deodara* (Roxb.) G.Don (Himalayan cedar) DNA. Reaction sets also included both positive (i.e., 2 ng total DNA from cultured *P. agathidicida* isolates NZFS 3128 or ICMP 18244) and negative (i.e., no DNA) controls.

We also evaluated assay sensitivity in terms of target copy number. As a template for these tests, we produced a PCR fragment 799 base pairs (bp) in length. Amplifications were typically performed in 20 μL reaction volumes containing 1× EmeraldAmp GT PCR Master Mix (Takara Bio Inc., Kusatsu City, Shiga Prefecture, Japan) and 0.5 μM of each amplification primer (PTA_pcrF, 5’-CCAAACATAGCTATAACCCCACCA-3’; PTA_pcrR, 5’-GGTTTCGGTTCGTTAGCCG-3’). Thermocycling was performed using a T1 Thermocycler (Biometra GmbH, Göttingen, Germany) and with standard cycling conditions including an initial 4 min denaturation at 94°C, then 35 cycles of 94°C for 30 secs, 58°C for 30 secs and 72°C for 30 secs, with a final 5 min extension at 72°C. Amplification products were prepared for DNA sequencing using shrimp alkaline phosphatase (ThermoFisher Scientific, Waltham, MA) and exonuclease 1 (ThermoFisher Scientific), following a manufacturer recommended protocol. Sensitivity tests using the PCR fragment as template were conducted as for total DNA.

### Comparison of standard bioassay and hybrid LAMP bioassay

A direct comparison of the extended bioassay and hybrid LAMP assay was performed for soil samples collected from sites in the Waitākere Ranges Regional Park and in the Waipoua Forest Sanctuary (S2 Table). Samples from each site typically consisted of 1-2 kg of soil from the upper 15 cm of the mineral horizon in the vicinity of kauri trees displaying dieback symptoms. Subsamples of 500 g were first air dried in open plastic containers for two days, moist incubated for four and subsequently flooded with 500 mL of reverse osmosis H_2_O. Fifteen detached *Cedrus deodara* needles were then floated on the water surface. Baits were removed after 48 h (Fig 1); ten were immediately used for the standard bioassay with the remainder frozen at −20°C prior to DNA extraction and LAMP testing.

**Fig 1.**
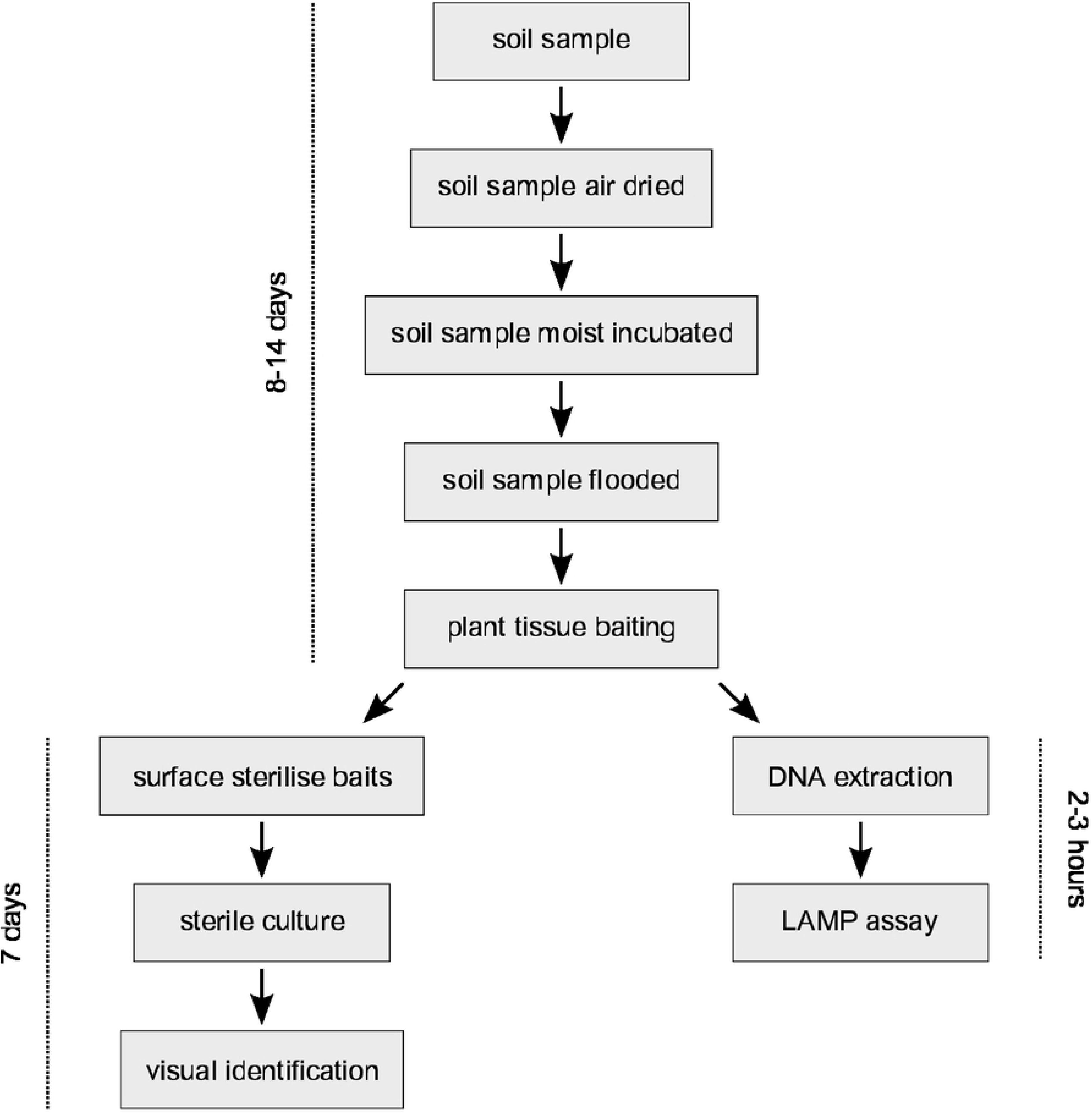
Schematic diagram comparing the extended bioassay and hybrid LAMP assay workflows and timelines.

For the standard bioassay, cedar baits were first rinsed with reverse osmosis (RO) H_2_O, soaked in 70% ethanol for 30 s, then rinsed again with RO H_2_O before being dried on clean paper towels. Surface sterilised baits were then placed on *Phytophthora-selective* media [37] using sterile technique and incubated at 18°C for 5-7 days. To identify the resulting cultures, asexual and sexual structures were examined using a Nikon ECLIPSE 80i compound light microscope (Nikon Corporation, Tokyo, Japan) with micrographs captured using a Nikon DS-Fi1 digital microscope camera head (Nikon Corporation) and processed using NIS-Elements BR (version 5.05, Nikon Corporation) [8, 38] (Fig 1).

For the hybrid LAMP assay total DNA was extracted from two or three frozen cedar baits using the Machery-Nagel Plant Kit II (Macherey-Nagel GmbH & Co. KG, Düren, Germany) and the manufacturer’s recommended protocol for plant material. Following extraction, the concentration of total DNA was determined for each sample using a Qubit 2.0 Fluorometer (Thermo Fisher Scientific, Waltham, MA). LAMP assays were performed using optimised reaction mixes and conditions with up to 5 ng of total bait DNA added as template. Each reaction set included both positive (i.e., 2 ng total DNA from a cultured isolate of *P. agathidicida*) and negative (i.e., milliQ H_2_O) controls (Fig 1).

### Sequencing of total bait DNA

We used whole genome sequencing of the DNA extracted from cedar baits from three sites in the Waitākere Ranges Regional Park to evaluate results from the extended and hybrid bioassays. Shotgun sequencing libraries were prepared for each DNA extraction using Illumina Nextera DNA library preparation kits (Illumina, Inc., San Diego, CA). The Massey Genome Service (Palmerston North, New Zealand) performed library preparation, paired-end DNA sequencing and quality assessment of the resulting reads.

For each sample, we performed a preliminary *de novo* assembly using idba_ud [39]. The resulting contigs were then compared to our collection of mitochondrial genome sequences (S1 Table) using BLAST [36]. Using the reference assembly tool implemented in Geneious R9 [35], contigs with high similarity to the reference set were then mapped to the mitochondrial genome sequences of *P. agathidicida* (NZFS 3118; Winkworth et al., unpubl.), *P. cinnamomi* (NZFS 3750; Winkworth et al., unpubl.), and *Pythium ultimum* [40]. Assemblies were subsequently checked by eye; contigs were removed from an assembly if similarity to another reference genome was higher.

## Results

### Target region identification and primer design

A comparison of mitochondrial genome sequences for members of *Phytophthora* clade 5 suggested several potential targets for a LAMP assay specific to *P. agathidicida*. Seven sets of LAMP primers, each consisting of between four and six primers, were designed using PrimerExplorer v5. However, in initial trials (data not shown) all but one of these primer sets failed to discriminate *P. agathidicida* from other Clade 5 species. We assume that although there are sequence level differences between *P. agathidicida* and other Peronosporaceae at these loci, the overall number and/or distribution of these differences is not sufficient to prevent amplification in non-target species. The exception was a set of six primers targeting a 227 nucleotide long section of the apocytochrome b (*cob*) coding sequence spanning from nucleotide position 392 to position 617 (Table 1; Fig 2). Of the initial primer sets, only this one was tested further.

**Fig 2.**
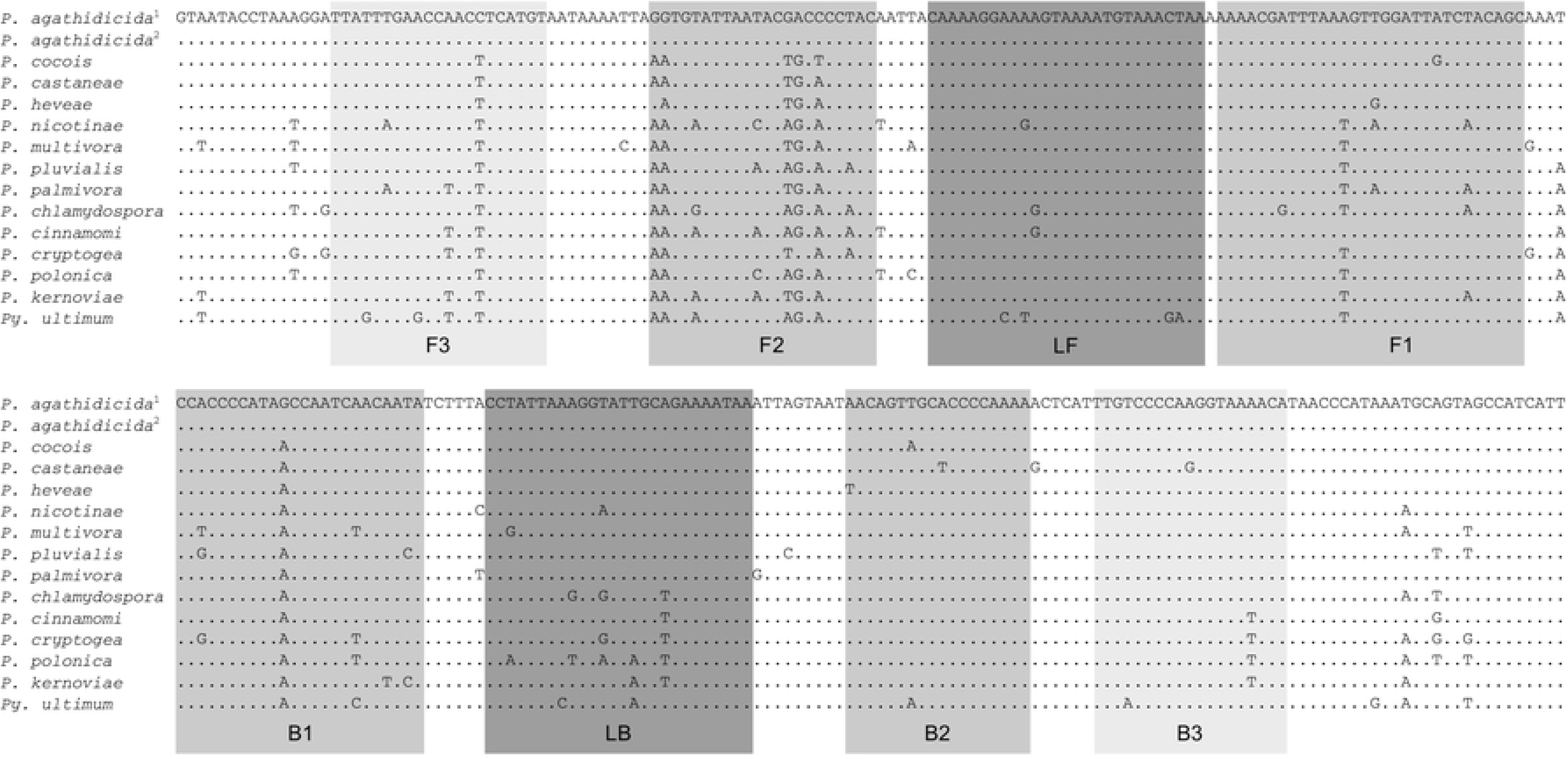
Multiple sequence alignment for 13 representative *Phytophthora* species plus *Pythium ultimum* for a section of the mitochondrial genome containing the *P. agathidicida* LAMP assay target. LAMP primer binding sites are indicated by grey outlines; F3 and B3 are binding sites for the external primer pair (i.e., PTAF3 and PTAB3), F2/F1 and B2/B1 form the binding sites for the internal primers (i.e., PTAFIP and PTABIP, respectively), and LF and LB are binding sites for the loop primers (i.e., PTALAF and PTALB)

**Table 1.**
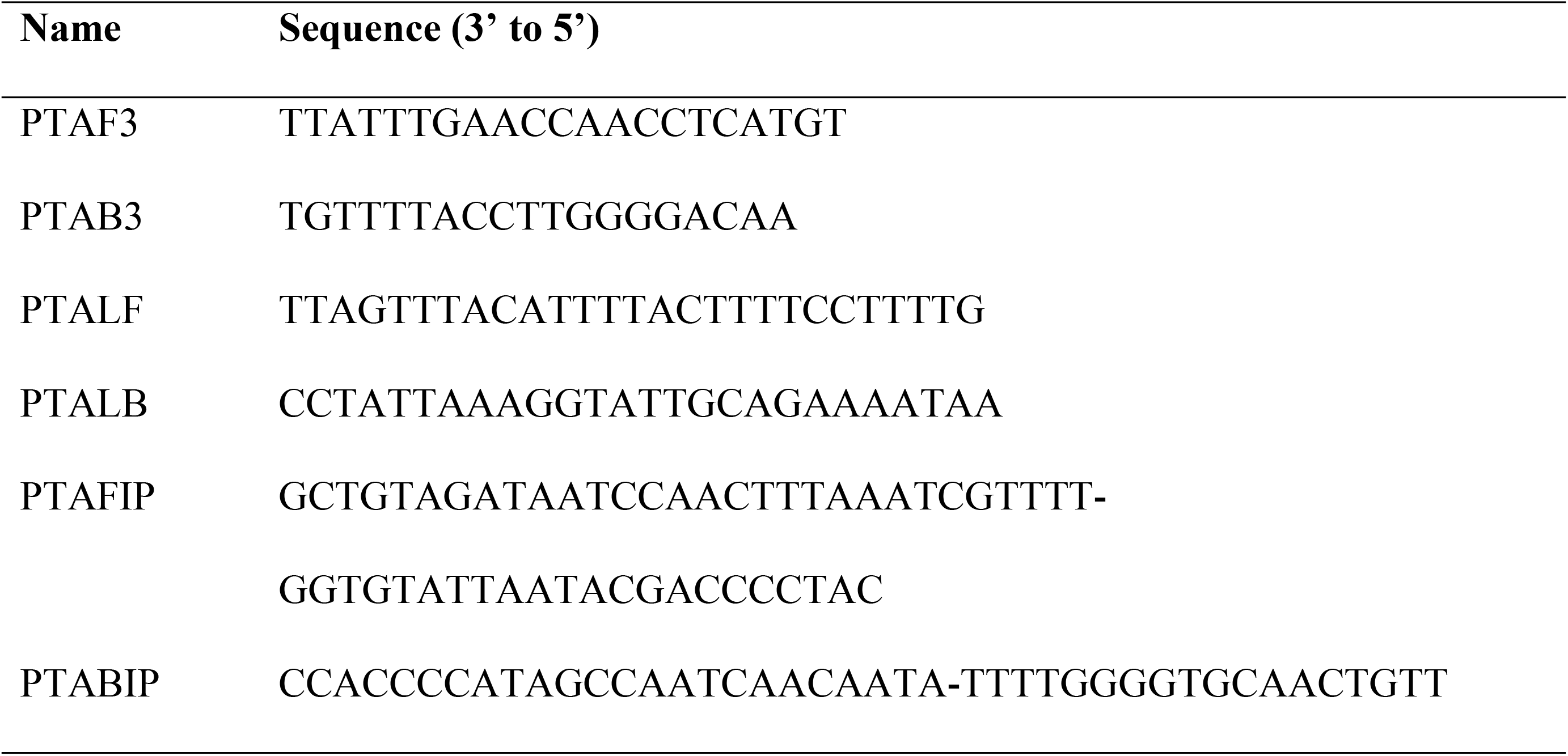
Primer sequences for the *P. agathidicida* loop-mediated isothermal amplification (LAMP) assay.

### LAMP assay optimisation

Using 2 ng of total DNA from cultured *P. agathidicida* isolates NZFS 3128 or ICMP 18244 as template, the *P. agathidicida* LAMP assay gave similar results across a range of reaction conditions. Specifically, amplification was observed for all examined ratios of external to internal primers (i.e., 1:3, 1:4, 1:6 and 1:8), amplification temperatures (i.e., 60°C, 63°C, and 65°C), and amplification times (i.e., 30 min, 45 min, 60 min, and 90 min). Conversely, no amplification was observed under these same reaction conditions for controls containing no DNA.

For subsequent analyses, a 1:3 ratio of external to internal primers, an amplification temperature of 63°C, and an amplification time of 45 min were used.

### LAMP assay specificity and sensitivity

Using optimised reaction mixes and conditions, we consistently recovered amplification products from tested *P. agathidicida* isolates (Table 2). In all cases, amplification was detected using real-time fluorescence (e.g., Fig 3, panel B curves B and C) and agarose gel electrophoresis (e.g., Fig 3, panel A lanes B and C). Conversely, amplification was not detected using either agarose gel electrophoresis or real-time fluorescence for reactions containing no DNA (e.g., Fig 3, panel A curve H and panel B lane H) or those containing total DNA from other representatives of *Phytophthora* (Table 2).

**Fig 3.**
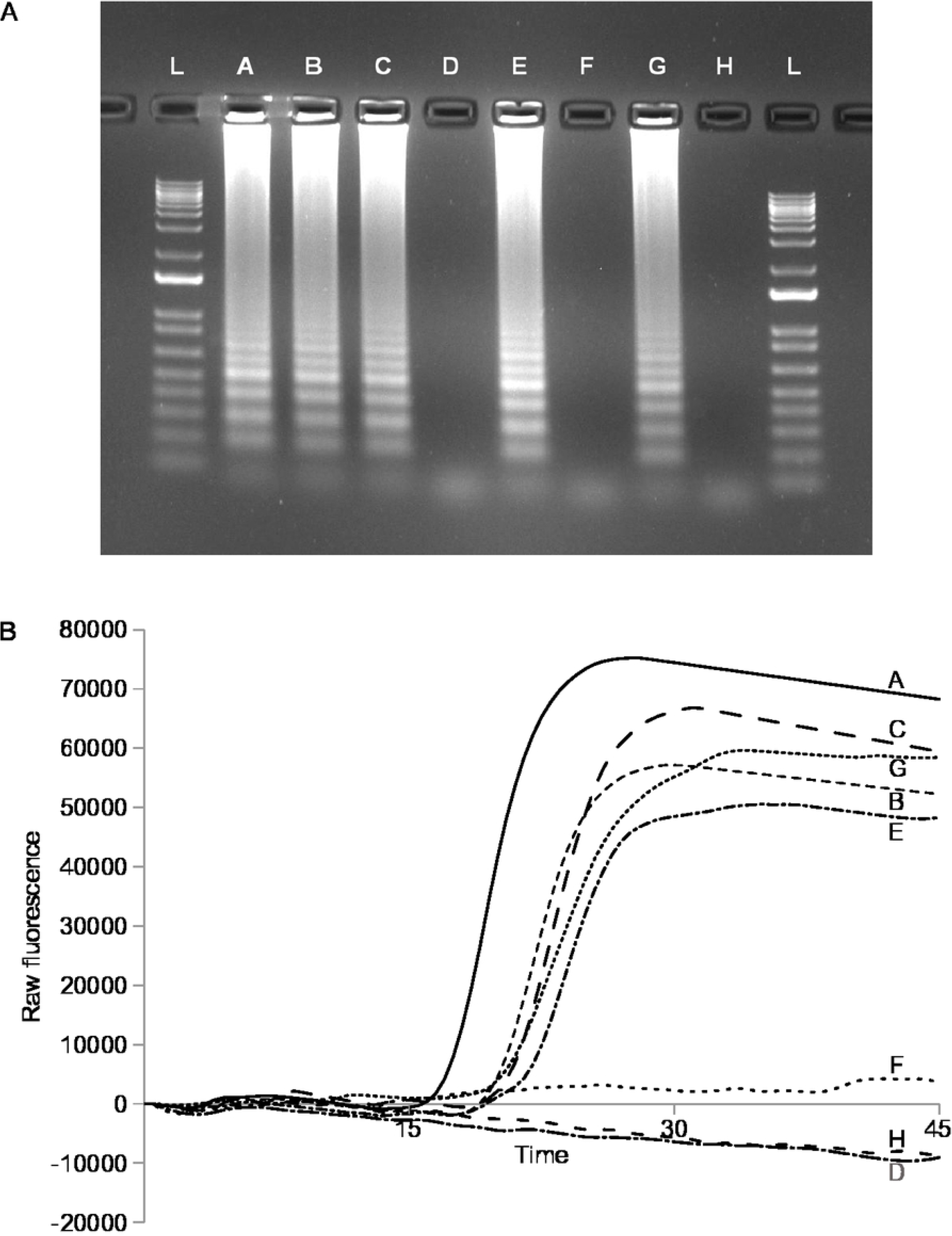
Results of the *P. agathidicida* LAMP assay. A. Endpoint visualisation of LAMP products using SYBR Safe (Invitrogen) following electrophoresis on a 1% TAE agarose gel. Lane A, 2 pg PCR products; lane B, 2 pg total DNA isolate ICMP18244; lane C, 2 pg total DNA isolate ICMP18410; lane D, 5 ng total DNA from baited Waitakere sample 1018; lane E, 5 ng total DNA from baited Waitakere sample 1020; lane F, 5 ng total DNA from baited Waipoua Forest sample 1072; lane G, 5 ng total DNA from baited Waipoua Forest sample 1081; lane H, no DNA control; lane L, 1 kb plus DNA ladder. B. Real-time visualisation of LAMP products using raw fluorescence data. Samples labelled as for A.

**Table 2.**
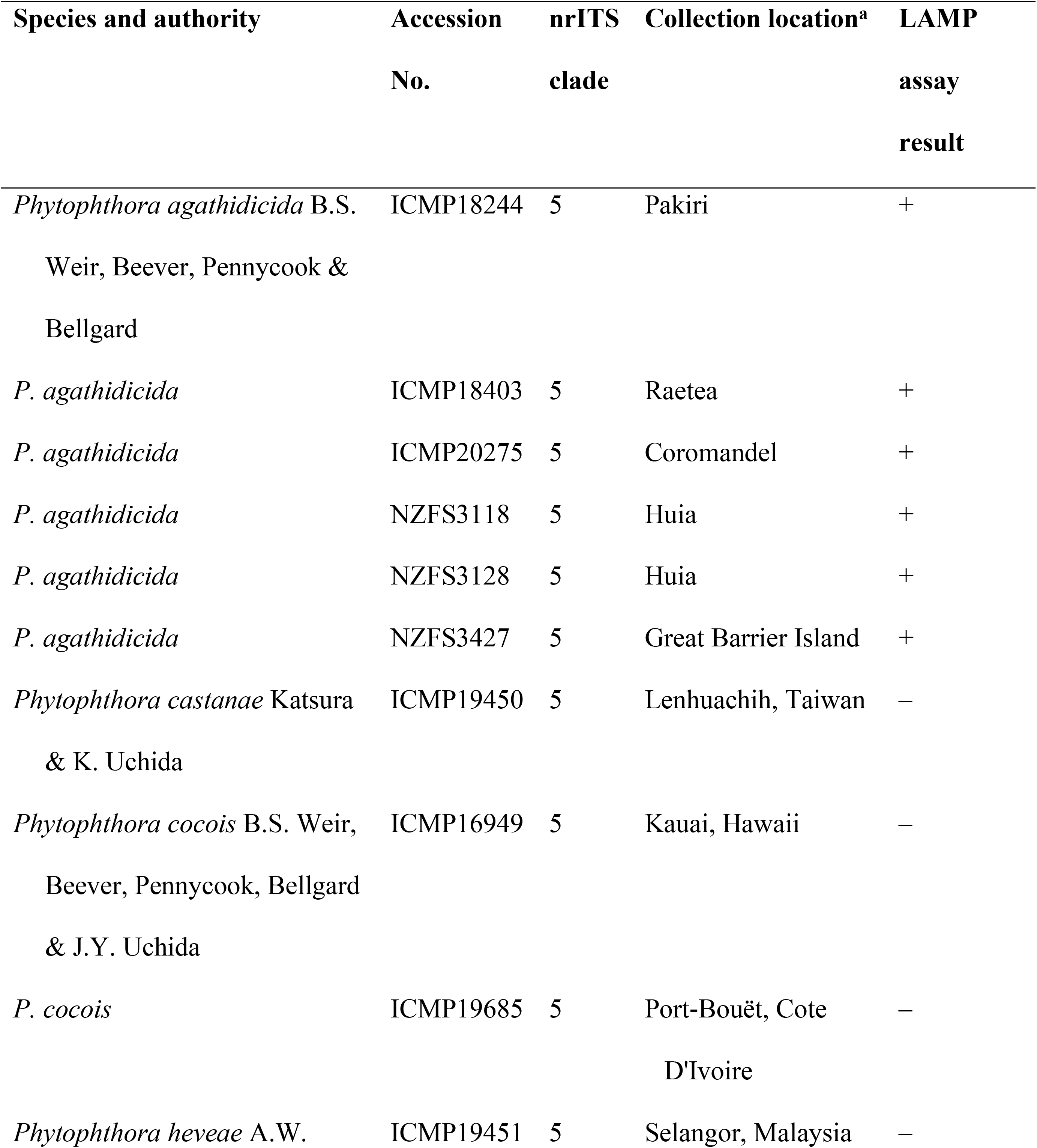
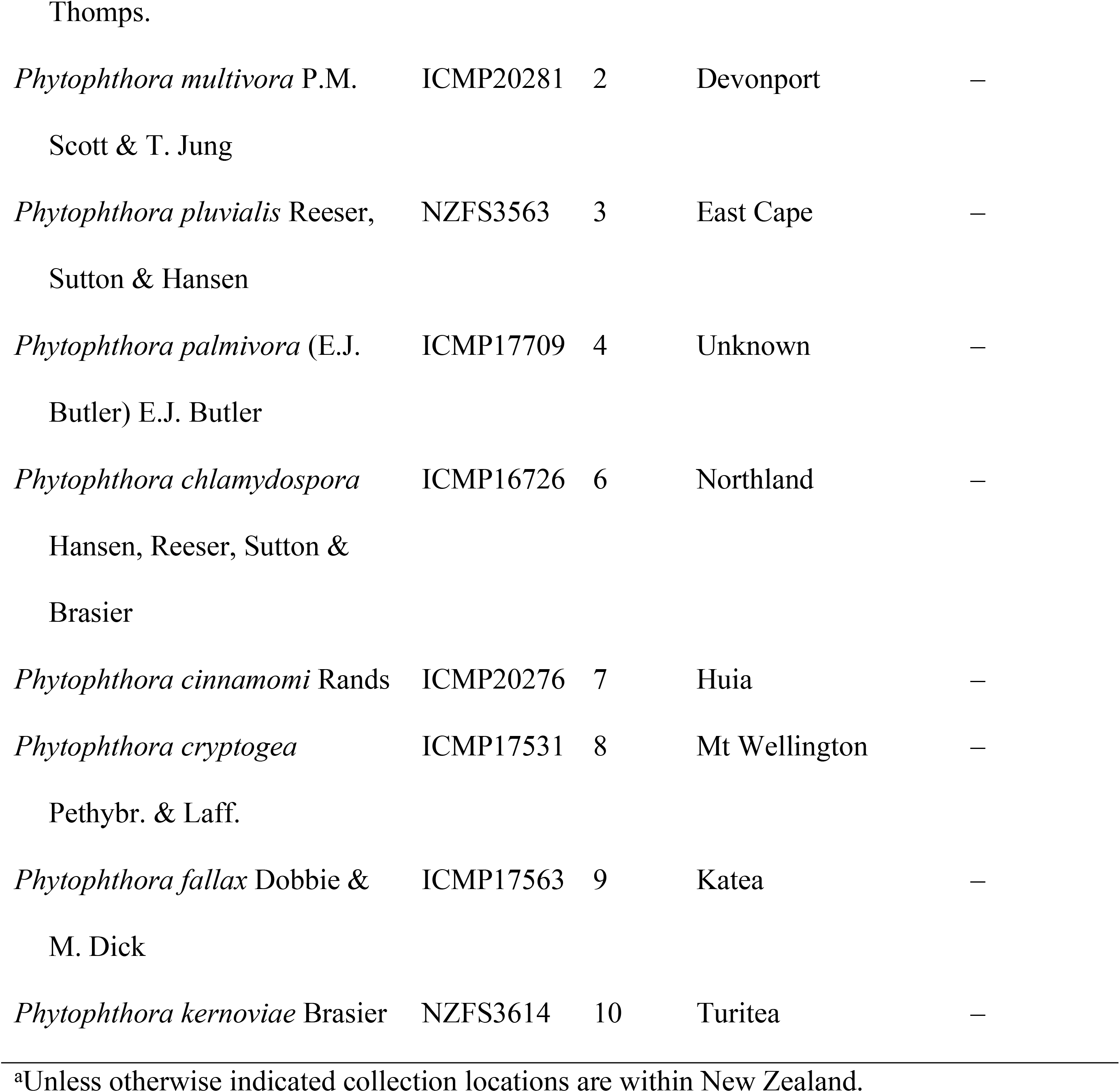
*Phytophthora* isolates used in specificity testing and results of testing with the *P. agathidicida* LAMP assay for these isolates.

We initially examined assay sensitivity using total *P. agathidicida* DNA. Using optimised reaction mixes and conditions, we consistently detected as little as 1 fg total DNA from cultured *P. agathidicida* isolate ICMP 18244; this limit remained the same when 2 ng total *Cedrus deodara* DNA was also added to LAMP reactions. The detection limit when using a PCR amplification product containing the target locus as template was 100 ag. Again, this limit was unchanged by the addition of 2 ng total *Cedrus deodara* DNA. Given Avogadro’s number (i.e., 6.022 × 10^23^ molecules/mole), the predicted length of the amplification product (i.e., 799 bp), and average weight of a base pair (i.e., 650 Daltons) the observed detection limit of 100 ag corresponds to 116 copies of the target fragment.

### Comparison of standard bioassay and hybrid LAMP bioassay

The extended and hybrid LAMP bioassays produced contrasting results for soil samples from sites in the Waitākere Ranges Regional Park and Waipoua Forest Sanctuary (Table 3). Using the extended bioassay, *P. agathidicida* was detected in one of six soil samples from the Waitākere Ranges Regional Park and none of the six from the Waipoua Forest Sanctuary. Detections were five out of six for the Waitākere Ranges Regional Park and three of six for the Waipoua Forest Sanctuary using the LAMP assay to test DNA extracted from cedar baits.

**Table 3.** Comparison of *Phytophthora agathidicida* detections from field collected soil samples using the extended bioassay and *P. agathidicida* loop-mediated isothermal assay.

| Accession no.<sup>a</sup> | Collection location | Extended bioassay result | Other species detected by the extended bioassay | LAMP assay result |
| --- | --- | --- | --- | --- |
| Waitākere Ranges Regional Park |  |  |  |  |
| HTHF1003 | Maungaroa Ridge | – | *P. cinnamomi*,<br>*Pythium* sp. | + |
| HTHF1014 | Maungaroa Ridge | – | – | + |
| HTHF1018 | vicinity of Lower Kauri Track | – | *P. cinnamomi*,<br>*Pythium* sp. | – |
| HTHF1020 | vicinity of Lower Kauri | + | – | + |

|  | Track |  |  |  |
| --- | --- | --- | --- | --- |
| HTHF1035 | vicinity of Huia Dam | + | – | + |
| HTHF1037 | vicinity of Huia Dam | – | *P. cinnamomi*,<br>*Pythium* sp. | + |
| Waipoua Forest Sanctuary |  |  |  |  |
| HTHF1043 | State Highway 12 | – | – | + |
| HTHF1055 | State Highway 12 | – | *P. cinnamomi* | – |
| HTHF1081 | State Highway 12 | – | *Pythium* sp. | + |
| HTHF1082 | State Highway 12 | – | – | + |
| HTHF1091 | Waipoua River Bridge | – | – | – |
| HTHF1092 | Waipoua River Bridge | – | *P. cinnamomi* | – |

<sup>a</sup>Samples drawn from a wider set collected as part of the “Healthy Trees Healthy Futures” programme.

Mitochondrial genome sequences assignable to *P. agathidicida*, *P. cinnamomi* and a member of the genus *Pythium* were recovered from whole genome sequencing of cedar bait DNA from three sites in the Waitākere Ranges Regional Park. In the case of *P. agathidicida*,contigs corresponding to 98.6-100% of the reference mitochondrial genome were recovered; smaller segments of the representative of the *P. cinnamomi* and *Pythium* genomes were recovered. Recovery of *P. agathidicida* genomes is consistent with results of the LAMP assay while recovery of sequences from other oomycetes is consistent with these lineages being detected on the basis of morphology following culturing.

## Discussion

Molecular assays have been reported for various *Phytophthora* species [e.g., 28, 41, 42]. In the present study, we have developed a LAMP assay for the detection of *P. agathidicida*, the causative agent of kauri dieback. When combined with soil baiting, this assay, which targets a region of the mitochondrial apocytochrome b gene, provides a powerful alternative to the currently used extended soil bioassay.

In specificity testing, our LAMP assay did not cross react with a range of *Phytophthora* species, including all recognised members of Clade 5 and four of the remaining five species from kauri forest soils (Table 3). These results are consistent with pairwise comparisons of the assay target sequence from *P. agathidicida* and 64 other Oomycete taxa (S3 Table). These comparisons indicate pairwise sequence differences of 2.6-15.5% across the entire set; the other Clade 5 species differed by 2.6-3.7% and remaining kauri forest species by 5.8.0-9.6 %. Together, these results suggest our LAMP assay provides a specific test for the presence of *P. agathidicida*.

Our LAMP assay consistently detected 1 fg total *P. agathidicida* DNA. This limit is lower than that of several other *Phytophthora-specific* LAMP assays [e.g., 29] but similar to the limit reported by Than et al. [15] for their *P. agathidicida* RT-PCR assay (i.e., 2 fg). However, unlike the Than et al. [15] assay, which was ten-fold less sensitive in the presence of soil DNA, in our testing the sensitivity of the LAMP assay was unchanged in the presence of background DNA. This result suggests that our LAMP assay is likely to outperform the Than et al. [15] assay for complex samples (e.g., soil or bait DNA). Although it is common to report assay detection limits in terms of the total amount of DNA, these can be difficult to interpret and we therefore also estimated a detection limit based on target copy number.

Using a PCR-amplified target fragment the observed detection limit was approximately 116 target copies. We are not aware of estimates for the numbers of mitochondrial genomes per cell in Oomycetes but for *Saccharomyces cerevisae* 20-200 mitochondrial genome copies per cell have been reported [43]. While we acknowledge there is considerable variation in mitochondrial genome counts (e.g., between taxa or life stages), the *S. cerevisae* count implies relatively few cells would need to have colonized a plant bait in order to detect *P.agathidicida*. Indeed, testing in this study suggests that the sensitivity of our LAMP assay is sufficient to detect *P. agathidicida* at the levels typically encountered on cedar baits used for soil baiting.

For samples from both the Waitakere Ranges Regional Park and Waipoua Forest Sanctuary, *P. agathidicida* was detected with higher frequency using the LAMP assay than the morphology-based approach typically used with the bioassay (Table 3). For example, *P. agathidicida* was detected in one of six Waitakere Ranges Regional Park samples based on visual inspections of cultures whereas five of these same samples were positive for *P. agathidicida* based on the LAMP assay (Table 3). Whole genome sequencing of DNA extracted from cedar baits supports LAMP results. That complete, or nearly so, mitochondrial genome sequences were recovered from each of the three samples examined argues strongly for *P. agathidicida* having colonised the baits, as does the observation that in preliminary phylogenetic analyses, these sequences were most closely related to others from the Waitakere Ranges. For example, the sequence from soil sample HTHF 1035 that was collected at Huia fell sister to two previously sequenced Huia samples (i.e., NZFS 3118 and NZFS 3128) in our analyses. Also consistent with visual inspections of cultures, our whole genome sequencing recovered sequences assignable to the mitochondrial genomes of *P. cinnamoni* and *Pythium* sp. (Table 3).

For six of the samples, *P. agathidicida* was not detected using the extended bioassay despite the LAMP assay and, for some samples, whole genome sequencing suggesting that zoospores of this species had colonised baits (Table 3). Indeed, this contrast implies that the extended bioassay, and more specifically culturing from bait tissues, may be systematically biased against the recovery of *P. agathidicida*. If so, the slow average *in vitro* growth rate of *P. agathidicida* (4.5 mm/day; [8]) may provide at least a partial explanation. Specifically, for three samples where the extended bioassay and LAMP testing results differed, oomycete species with faster *in vitro* growth rates – *P. cinnamomi* (8.3 mm/day [44]) and *Pythium* sp. (21-29 mm/day [45]) – were identified from cultures. Our ability to visually detect *P. agathidicida* may be limited by these other faster growing species; an issue that, assuming a standardised assay protocol, may not be overcome by parallel processing of split replicate soil samples in different laboratories. That said, both Beaver et al. [46] and McDougal et al. [47] have reported inconsistent recovery of *P. agathidicida* in such situations. However, the protocol used in these studies allowed for the use of different bait tissues (i.e., detached cedar needles or intact lupin radicals) and recently Khaliq et al. [48] have shown that bait type and integrity (e.g., detached or intact) can influence the recovery of *Phytophthora* by culturing. In either case the impacts of such biases are likely to be reduced by genetically testing the baits themselves.

Initially, we envisage using the LAMP assay to directly test bait tissues recovered from flooded soils. The use of baiting minimises the impacts of soil heterogeneity and low pathogen titre on subsequent detection [e.g., 28, 47] while use of the LAMP assay for detection increases the sensitivity and reproducibility of testing. This hybrid approach is also faster and more cost effective than the extended bioassay; the approach does away with the time and expense of culturing and initial morphological identification, as well as is the need for confirmatory sub-culturing and qPCR testing. Moreover, because culturing is not required, testing can be performed without the need for centralised laboratory facilities. Although in the present study we use a DNA extraction protocol suited to a laboratory situation, alternatives suitable for use outside a laboratory have also been successfully trialled. This testing approach has the potential to dramatically enhance our ability to address the threat posed by kauri dieback. For example, we could move beyond confirmatory testing to active monitoring of the pathogen across the distribution of kauri. This could provide a direct estimate of pathogen distribution, measures of rate and pattern of spread, and a means to evaluate the efficacy of disease control measures. This assay also has the potential to empower individual landowners and community groups in a way that has not previously been possible. Direct access to a diagnostic tool will allow them to evaluate pathogen status in their area, make immediate land management decisions, and engage in an informed way with regional and national initiatives.

## Supporting Information

**S1 Table. Sources of the oomycete mitochondrial genome sequences used when designing the *P. agathidicida* loop-mediated isothermal amplification (LAMP) assay.**

(DOC)

**S2 Table. Location details for field collected soil samples used in comparisons of extended bioassay and *P. agathidicida* loop-mediated isothermal assay.**

(DOC)

**S3 Table. Comparison of the *Phytophthora agathidicida* LAMP assay target sequence and the corresponding locus in other Oomycetes.**

(DOC)

## Acknowledgements

We acknowledge the mana whenua Te Kawerau ā Maki (Waitākere Ranges) and Te Roroa (Waipoua). We thank the International Collection of Microorganisms from Plants (ICMP) and New Zealand Forest Research Institute (NZFS) culture collections as well as the Healthy Trees Healthy Forests (HTHF) program for access to samples. We also acknowledge financial support from the BioProtection Research Centre (P.J.L. and R.C.W.), Massey University (R.C.W. and P.J.L.), and the New Zealand Ministry of Business, Innovation and Employment via the Catalyst: Seeding Fund (administered by the Royal Society Te Apārangi; P.J.L. and R.C.W) and the Strategic Science Investment Fund (S.E.B).

## Author Contributions

**Conceptualization:** Richard C. Winkworth, Peter J. Lockhart.

**Formal Analysis:** Richard C. Winkworth.

**Investigation:** Richard C. Winkworth, Briana C.W. Nelson, Chantal M. Probst, Stanley E. Bellgard, Patricia A. McLenachan.

**Methodology:** Richard C. Winkworth, Stanley E. Bellgard.

**Project Administration:** Richard C. Winkworth.

**Resources:** Stanley E. Bellgard, Chantal M. Probst.

**Validation:** Richard C. Winkworth, Briana C.W. Nelson.

**Visualization:** Richard C. Winkworth.

**Writing – Original Draft Preparation:** Richard C. Winkworth, Briana C.W. Nelson.

**Writing – Review & Editing:** Richard C. Winkworth, Briana C.W. Nelson, Stanley E. Bellgard, Chantal M. Probst, Patricia A. McLenachan, Peter J. Lockhart.

## References

1. Beever RE, Coffey MD, Ramsfield TD, Dick M, Horner IJ. Kauri (*Agathis australis*) under threat from *Phytophthora*? Fourth Meeting of IUFRO Working Party S07.02.09. General technical report PSW-GTR-221, pp. 74–85. U.S. Department of Agriculture, Forest Service Pacific Southwest Research Station, Monterey, CA.

2. Gadgil PD. *Phytophthora heveae*, a pathogen of kauri. New Zealand Journal of Forestry Science. 1974; 4: 59–63.

3. Waipara N, Hill S, Hill L, Hough E, Horner I. Surveillance methods to determine tree health, distribution of kauri dieback disease and associated pathogens. New Zealand Plant Protection. 2013; 66: 235–41.

4. Blair JE, Coffey MD, Park SY, Geiser DM, Kang SA. multi-locus phylogeny for *Phytophthora* utilizing markers derived from complete genome sequences. Fungal Genetics and Biology. 2008; 45, 266–77 https://doi.org/10.1016/j.fgb.2007.10.010.

5. Martin FN, Blair JE, Coffey MD. A combined mitochondrial and nuclear multilocus phylogeny of the genus *Phytophthora*. Fungal Genetics and Biology. 2014; 66: 19–32. https://doi.org/10.1016/j.fgb.2014.02.006.

6. McArthy CGP, Fitzpatrick DA. Phylogenomic reconstruction of the Oomycete phylogeny derived from 37 genomes. mSphere. 2017; 2:e00095–17. https://doi.org/10.1128/mSphere.00095-17.

7. Bourret TB, Choudhury RA, Mehl HK, Blomquist CL, McRoberts N, Rizzo DM. Multiple origins of downy mildews and mitonuclear discordance within the paraphyletic genus *Phytophthora*. PLOS ONE. 2018; 13: e0192502. https://doi.org/10.1371/journal.pone.0192502.

8. Weir BS, Paderes EP, Anand N, Uchida JY, Pennycook SR, Bellgard SE, et al. A taxonomic revision of *Phytophthora* Clade 5 including two new species, *Phytophthora agathidicida* and *P*. *cocois*. Phytotaxa. 2015; 205: 21–38.

9. Jeffers SN, Martin SB. Comparison of two media selective for *Phytophthora* and *Pythium* species. Plant Disease. 1986; 70: 1038–43.

10. Kato S, Coe R, New L, Dick MW. Sensitivities of various Oomycetes to hymexazol and metalaxyl. Journal of General Microbiology. 1990; 136: 2127–34

11. Newhook FJ. The association of *Phytophthora* spp. with mortality of *Pinus radiata* and other conifers. New Zealand Journal of Agricultural Research. 1959; 2: 808–43.

12. Martin FN, Abad ZG, Balci Y, Ivors K. Identification and detection of *Phytophthora:* reviewing our progress, identifying our needs. Plant Disease. 2012; 96: 1080–103.

13. Horner IJ, Wilcox WF. Temporal changes in activity and dormant spore population of *Phytophthora cactorum* in New York apple orchards. Phytopathology. 1996; 86: 1133–9.

14. Bellgard SE, Pennycook SR, Weir BS, Ho W, Waipara NW *Phytophthora agathidicida*. Forest Phytophthoras. 2016; doi:10.5399/osu/fp.5.1.3748.

15. Than DJ, Hughes KJD, Boonhan N, Tomlinson JA, Woodhall JW, Bellgard SE. A TaqMan real-time PCR assay for the detection of *Phytophthora* ‘taxon Agathis’ in soil, pathogen of Kauri in New Zealand. Forest Pathology. 2013; 43: 324–30.

16. Vashist SK. Point-of-Care Diagnostics: Recent Advances and Trends. Biosensors. 2107; 7: 62. doi:10.3390/bios7040062.

17. Manessis G, Gelasakis AI, Bossis I The challenge of introducing point of care diagnostics in farm animal health management. Biomedical Journal of Scientific & Technical Research. 2019; 14: BJSTR. MS.ID.002601.

18. Miles TD, Martin FN, Coffey MD. Development of rapid isothermal amplification assays for detection of *Phytophthora* spp. in plant tissue. Phytopathology. 2015; 105: 265–78.

19. Cevallos W, Fernández-Soto P, Calvopiña M, Fontecha-Cuenca C, Sugiyama H, Sato H, et al. LAMPhimerus: A novel LAMP assay for detecting *Amphimerus sp*. DNA in human stool samples. PLOS Neglected Tropical Diseases. 2017; 11: e0005672. https://doi.org/10.1371/journal.pntd.0005672.

20. Seki M, Kilgore PE, Kim EJ, Ohnishi M, Hayakawa S, Kim DW. Loop-mediated isothermal amplification methods for diagnosis of bacterial meningitis. Frontiers in Pediatrics. 2018; 6: 57. doi: 10.3389/fped.2018.00057.

21. Zhang SY, Dai DJ, Wang HD, Zhang CQ. One-step loop-mediated isothermal amplification (LAMP) for the rapid and sensitive detection of *Fusarium fujikuroi* in bakanae disease through NRPS31, an important gene in the gibberellic acid biosynthesis. Scientific Reports. 2019;9, Article number: 3726.

22. Aglietti C, Luchi N, Pepori AL, Bartolini P, Pecori F, Raio A, et al. Real-time loop-mediated isothermal amplification: an early-warning tool for quarantine plant pathogen detection. AMB Express. 2019; 9:50 https://doi.org/10.1186/s13568-019-0774-9.

23. Farooq U, Latif A, Irshad H, Ullah A, Zahur AB, Naeem K, et al. Loop-mediated isothermal amplification (RT-LAMP): a new approach for the detection of foot-and-mouth disease virus and its sero-types in Pakistan. Iranian Journal of Veterinary Research. 2015; 16: 331–33.

24. Best N, Rodoni B, Rawlin G, Beddoe T. The development and deployment of a field based loop mediated isothermal amplification assay for virulent *Dichelobacter nodosus* detection on Australian sheep. PLOS ONE. 2018; 13: e0204310. https://doi.org/10.1371/journal.pone.0204310.

25. Li Y, Fan P, Zhou S, Zang L. Loop-mediated isothermal amplification (LAMP): A novel rapid detection platform for pathogens. Microbial Pathogenesis. 2017; 107: 54–61.

26. Notomi T, Mori Y, Tomita N, Kanda H. Loop-mediated isothermal amplification (LAMP): principle, features and future prospects. Journal of Microbiology. 2015; 53: 1–5.

27. Dong ZM, Liu PQ, Li BJ, Chen GL, Weng QY, Chen QH. Loop-mediated isothermal amplification assay for sensitive and rapid detection of *Phytophthora capsici*. Canadian Journal of Plant Pathology. 2015; 37, 485–94.

28. Dai TT, Yang X, Hu T, Li ZY, Xu Y, Lu CC. A novel LAMP assay for the detection of *Phytophthora cinnamomi* utilizing a new target gene identified from genome sequences. Plant Disease. 2019; doi: 10.1094/PDIS-04-19-0781-RE.

29. Hansen ZR, Knaus BJ, Tabima JF, Press CM, Judelson HS, Grunwald NJ, et al. Loop-mediated isothermal amplification for detection of the tomato and potato late blight pathogen, *Phytophthora infestans*. Journal of Applied Microbiology. 2016; 120: 1010–20.

30. Khan MR, Li BJ, Jiang Y, Weng QY, Chen QH. Evaluation of different PCR-based assays and LAMP method for rapid detection of *Phytophthora infestans* by targeting the *Ypt1* gene. Frontiers in Microbiology. 2017; 8: 1920. https://doi.org/10.3389/fmicb.2017.01920.

31. Chen Q, Li B, Liu P, Lan C, Zhan Z, Weng Q. Development and evaluation of specific PCR and LAMP assays for the rapid detection of *Phytophthora melonis*. European Journal of Plant Pathology. 2013; 137: 597–607.

32. Li B, Liu P, Xie S, Yin R, Weng Q, Chen Q. Specific and sensitive detection of *Phytophthora nicotianae* by nested PCR and loop-mediated isothermal amplification assays. Journal of Phytopathology 2015; 163: 185–93.

33. Zhao W, Wang T, Qi RD. *Ypt1* gene-based detection of *Phytophthora sojae* in a loop-mediated isothermal amplification assay. Journal of Plant Diseases and Protection. 2015; 122: 63–73.

34. Edgar RC. MUSCLE: multiple sequence alignment with high accuracy and high throughput. Nucleic Acids Research. 2004; 32: 1792–7.

35. Kearse M, Moir R, Wilson A, Stones-Havas S, Cheung M, Sturrock S, et al. Geneious Basic: An integrated and extendable desktop software platform for the organization and analysis of sequence data. Bioinformatics. 2012; 28: 1647–9. https://doi.org/10.1093/bioinformatics/bts199.

36. Altschul SF, Madden TL, Schäffer AA, Zhang J, Zhang Z, Miller W, et al. Gapped BLAST and PSI-BLAST: a new generation of protein database search programs. Nucleic Acids Research. 1997: 25: 3389–402.

37. Jeffers SN. Identifying species of Phytophthora. Clemson University. 2006. Available from: https://fhm.fs.fed.us/sp/sod/misc/culturing_species_phytophthora.pdf

38. Bellgard SE, Padamsee M, Probst CM, Lebel T, Williams SE. Visualising the early infection of *Agathis australis* by *Phytophthora agathidicida*, using microscopy and fluorescent in situ hybridizatisation assay. Forest Pathology. 2016; 46: 622–31 doi: 10.1111/efp.12280.

39. Peng Y, Leung HCM, Yiu SM, Chin FYL. IDBA-UD: a *de novo* assembler for singlecell and metagenomic sequencing data with highly uneven depth. Bioinformatics (Oxford, England). 2012; 28: 1420–8 https://doi.org/10.1093/bioinformatics/bts174.

40. Lévesque CA, Brouwer H, Cano L, Hamilton JP, Holt C, Huitema E, et al. Genome sequence of the necrotrophic plant pathogen *Pythium ultimum* reveals original pathogenicity mechanisms and effector repertoire. Genome Biology. 2010; 11: R73. doi: 10.1186/gb-2010-11-7-r73.

41. Schena L, Hughes KJD, Cooke DEL. Detection and quantification of *Phytophthora ramorum, P. kernoviae, P. citricola*, and *P. quercina* in symptomatic leaves by multiplex real-time PCR. Molecular Plant Pathology. 2006; 7: 365–79.

42. Minerdi D, Moretti M, Li Y, Gaggero L, Garibaldi A, Guillino ML. Conventional PCR and real time quantitative PCR detection of *Phytophthora cryptogea* on *Gebera jamesonii*. European Journal of Plant Pathology. 2008; 122: 227–37.

43. Williamson D. The curious history of yeast mitochondrial DNA. Nature Reviews Genetics. 2002; 3: 475–81.

44. Huberli D. Analysis of variability among isolates of *Phytophthora cinnamomi* Rands from *Eucalyptus marginata* Donn ex Sm. and *E. calophylla* R. Br. based on cultural characteristics, sporangia and gametangia morphology, and pathogenicity. Honours thesis, Murdoch University. 1995. Available from: https://researchrepository.murdoch.edu.au/id/eprint/1321/2/HUBERLI_whole_hons_thesis.pdf

45. Guo LY, Ko WH. Two widely accessible media for growth and reproduction of *Phytophthora* and *Pythium* species. Applied and Environmental Microbiology. 1993; 59: 2323–32.

46. Beever RE, Bellgard SE, Dick MS, Horner IJ, Ramsfield TD. Detection of *Phytophthora* taxon Agathis (PTA): Final Report. Landcare Report LC0910/137. 2010. Available from: https://www.kauridieback.co.nz/media/1640/11213-11215-12093-detection-of-phytophthora-taxon-agathis-pta-beever.pdf

47. McDougal R, Bellgard S, Scott P, Ganley B. Comparison of a real-time PCR assay and a soil bioassay technique for detection *Phytophthora* taxon Agathis from soil. Kauri Dieback Response, MPI Contract Report 53789. 2014. Available from: https://www.kauridieback.co.nz/media/1632/2014-17101-real-time-pcr-as-diagnostic-tool-comparison-of-a-real-time.pdf

48. Khaliq I, Hardy GES, White D, Burgess TI. 2018. eDNA from roots: a robust tool for determining Phytophthora communities in natural ecosystems. FEMS Microbiology Ecology. 2018; 94: fiy048

